# *Bacteroidales* on Harvesters: Baseline Prevalence and Abundance

**DOI:** 10.64898/2026.05.12.724369

**Authors:** Simerdeep Kaur, Jiangshan Wang, Ashley Kayabasi, Ishaan Rath, Ilan Benschikovski, Bibek Raut, Kyungyeon Ra, Mohit S. Verma

## Abstract

Fresh produce encounters pathogens at various stages of production and supply, with the harvesting process serving as one of these stages. To evaluate contamination associated with harvesting, we systematically swabbed zone 1 harvester surfaces and quantified *Bacteroidales* as a fecal biomarker using quantitative polymerase chain reaction (qPCR). Baseline contamination was dominated by non-detects, with occasional low-level detections (<25 copies/cm^2^) near the assay limit of detection (LoD). Detection occurred more frequently post-harvest (overall ∼4% pre-harvest and 10% post-harvest), while microbial loads remained low, indicating that harvesting primarily affected the likelihood of low-level contamination rather than increasing contamination abundance. Additionally, we developed and field-deployed a portable loop-mediated isothermal amplification (LAMP) assay for rapid harvester hygiene assessment and benchmarked its field performance against qPCR. Together, these results support a practical molecular tool for monitoring fecal contamination and informing cleaning and sanitization decisions.

## 1.1. Introduction

Contaminated harvesting equipment surfaces can transfer foodborne pathogens to fresh produce during harvesting processes, causing infections in humans upon consumption (Leaman et al., 2023). During the harvesting season, operators consistently use and relocate harvesting machines to various farms awaiting harvest (Leaman et al., 2023). Although harvesters are routinely cleaned after use, the rapid pace of operations necessitates fast verification of the cleaning process. Addressing this challenge involves two steps: first, identifying a specific biomarker to assess harvester hygiene and establishing its baseline levels; and second, ensuring its rapid detection to evaluate the potential risk posed by the equipment.

Harvester equipment’s produce contact surfaces and proximity areas are regions of concern. Commercial Food Sanitation classifies the surfaces into two zones: zone 1 includes produce contact surfaces that are easy to clean, while zone 2 lies near the produce and presents cleaning challenges (Leafy Greens Harvester Sanitation and Hygienic Design Working Group, 2021). Collaborative research between Commercial Food Sanitation and the Leafy Greens Marketing Agreement shows that Periodic Equipment Cleaning (PEC) effectively removes soil and microbial contaminants that tend to accumulate in zone 2. Designing and evaluating hygienic programs to mitigate these risks involves a thorough risk assessment before and during harvesting operations, employing microbiological and chemical testing methods.

Traditional lab-based microbiological technologies are impractical for cleaning verification, as they take at least a day to provide results. To facilitate uninterrupted equipment usage, quick cleaning verification methods are required. Adenosine triphosphate (ATP)-based assays enable rapid assessment of equipment cleanliness. These assays produce light (bioluminescence) when detecting ATP, an energy-carrying molecule present in living organisms. Despite their rapid results, a significant limitation is that ATP detected on equipment surfaces may come from soil, plant residues, or microorganisms (Corbitt et al., 2000; Mildenhall and Rankin, 2020). Some argue that defining cleanliness requires detecting organic debris beyond microbes (Davidson et al., 1999). However, the bacterial adherence can vary depending on the harvester’s design, material, and surface condition, making routine microbial level checks essential in such cases (Kleine et al., 2019; Verran et al., 2001). Furthermore, ATP testing lacks the capability to detect low levels of bacteria, requiring additional microbial techniques for confirmation (Lane et al., 2020; Leaman et al., 2023). Therefore, a test that can provide the specificity and sensitivity of microbial techniques and the speed of ATP-based assays would be an ideal candidate for evaluating harvester hygiene.

Our previous work shows that *Bacteroidales* serve as reliable indicators of fecal contamination (Wang et al., 2023). *Bacteroidales* are anaerobic bacteria and are commonly found in the mammalian gut. They are highly abundant, constituting 30-40% of total fecal bacteria, with a concentration of 10^9^ to 10^11^ colony-forming units (CFU) per gram of feces (Wang et al., 2024c). Importantly, their natural abundance from non-fecal sources is low. Given this range of abundance, *Bacteroidales* serve as a quantitative measure of fecal contamination. Unlike pathogenic detection, using *Bacteroidales* does not require enrichment before detection. In typical fresh produce harvesting areas distant from animal sources, *Bacteroidales* levels are low (0 (or <LoD) to 2 copies/cm^2^) (Wang et al., 2024c), while around animal operations, they are considerably higher (∼10^4^ copies/cm^2^) (Wang et al., 2023).

This study employs *Bacteroidales* to assess the variance in potential contamination across different sections of harvester zone 1 and times during operations. In 2022, we collected 400 swab samples from zone 1 of fresh-pack and processed lettuce harvesters in California. Samples were analyzed using quantitative polymerase chain reaction (qPCR) to characterize contamination patterns across harvester sections and harvest stages. Baseline contamination was dominated by non-detects, with detection occurring more frequently post-harvest than pre-harvest. When present, *Bacteroidales* concentrations were typically low (<25 copies/cm^2^), with two samples of elevated concentrations up to 93 copies/cm^2^. Furthermore, we used a colorimetric loop-mediated isothermal amplification (LAMP) integrated into microfluidic Paper-based Analytical Devices (µPADs) for on-site detection of *Bacteroidales* (Wang et al., 2024a). We deployed a portable heating and imaging system, Field-Applicable Rapid Microbial Loop-mediated Isothermal Amplification Platform (FARM-LAMP), to quantify fecal contamination using µPADs-based LAMP assay for evaluating harvester hygiene on farms (Wang et al., 2024a). With a reaction time of 60 minutes, this assay delivers cleaning verification of harvesting machines within one hour. This application paves the way for integrating nucleic acid amplification tests (NAATs) into standard practices for harvesting fresh produce, aiming to enhance food safety protocols.

## 2. Materials and Methods

### 2.1. Sample collection and processing

The research team traveled to California and collected a total of 204 swab samples from processed lettuce harvesters (Fig. S1) and 196 swab samples from fresh-pack lettuce harvesters (Fig. S2). All samples were shipped to Purdue University for downstream molecular analyses. For processed lettuce harvesters, swab samples were collected from multiple produce-contact and adjacent surfaces, including the conveyor belt, conveyor belt wall, curtain, tunnel, elevator, and funnel. Each surface was sampled over a defined 10 × 10 cm^2^ area. For fresh-pack lettuce harvesters, swab samples were collected from the plastic linings covering the packing tables. A 10 × 10 cm^2^ area of the packing table surface was swabbed both before and after harvesting (Fig. S2). To ensure consistent sampling area, a plastic template with a 100 cm^2^ square opening was placed over the surface prior to swabbing, and samples were collected directly from the exposed plastic lining. In all cases, BD CultureSwab™ Sterile Double Swabs (220135, BD, USA) were used for sample collection. Following swabbing, each swab pair was resuspended in 500 μL of nuclease-free water prior to molecular testing.

### 2.2. Quantitative polymerase chain reaction (qPCR)

qPCR assays were run on the swab resuspension samples, following protocols for the Luna® Universal Probe qPCR Master Mix (New England Biolabs, M3004). Each 20 µL reaction mix included 10 µL master mix, 0.8 µL PCR forward primer, 0.8 µL PCR reverse primer, 0.4 µL qPCR probe, 7 µL nuclease-free water, and 1 µL template. The qPCR primers and probe are shown in (Table S1). The assays were performed using PCR 96-plates (Catalog number AB0800W, Thermo Scientific) loaded in a qTOWER^3^ thermocycler (Analytik-Jena, Germany) with an initial denaturation of 95 °C for 60 s and a cycling profile of 95 °C for 15 s, 55 °C for 15 s, and 60 °C for 30 s. The results of the qPCR assays were compared with those obtained from FARM-LAMP.

### 2.3. Determining *Bacteroidales* levels from qPCR results

Using the quantification cycle (Cq) values of each sample, the log_10_(copies/µL) was calculated using the Equation 1 from the qPCR calibration reported by Wang *et al* (Wang et al., 2024c). The log_10_(copies/µL) values were then converted to copies/µL using Equation 2. Next, the number of copies/cm^2^ was calculated by multiplying the copies/µL by the resuspension volume of water (500 µL) and dividing the result by 100 (the area swabbed in cm^2^) (Equation 3). Finally, the log_10_(copies/cm^2^) was determined by taking the base-10 logarithm of the copies/cm^2^ value (Equation 4).

1. *log*_*10*_ *Copies per* 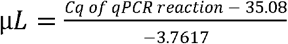
2. *Copies per* 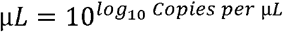
3. *Copies per* 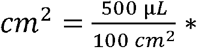 (*copies per reaction in* 1 *µL*)
4. *log*_*10*_ *Copies per cm*^*2*^ *= log*_*10*_ *= (Copies per cm*^*2*^)

### 2.4. Determining Limit of Quantification (LoQ) and Limit of Detection (LoD)

First, a standard curve was generated by ordinary least squares regression using the calibrator concentrations:

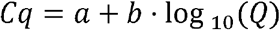

where *Cq* is cycle threshold and *Q* is copies per reaction. For each replicate Cq at a given concentration, the corresponding back-calculated quantity was obtained from the inverse of the standard curve:

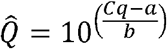

For each concentration, precision was calculated on the back-calculated quantities 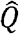 as:

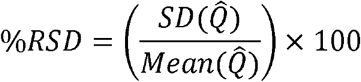

The LoQ was set as the lowest concentration with %RSD ≤ 35% among replicate reactions (Forootan et al., 2017). LoD was conservatively reported as the lowest tested concentration with 100% detection across replicates.

### 2.5. Analysis of qPCR results

qPCR data were analyzed to characterize both the prevalence and abundance of *Bacteroidales*. All analyses were conducted in Python using pandas, NumPy, statsmodels, and matplotlib. Reactions with missing Cq values were assigned a value of 45 (last qPCR cycle) and were classified as non-detects (ND). Detection thresholds were defined and applied uniformly across all datasets. The LoD was determined as 1 copy per reaction (equivalent to 5 copies/cm^2^), and the LoQ was determined as 5 copies per reaction (equivalent to 25 copies/cm^2^) (see section 2.4). Samples with detectable signals at or above the LoD but below the LoQ were classified as detectable but not quantifiable (DNQ), while samples at or above the LoQ were classified as detectable and quantifiable (DQ).

The qPCR dataset was characterized by a high proportion of NDs and sparse quantifiable observations, therefore, contamination dynamics were evaluated using a two-part (hurdle-type) analytical framework that separates contamination prevalence from contamination abundance. Contamination prevalence is defined as detected samples (DQ + DNQ)/total number of samples tested for the condition. Prevalence was reported with Wilson 95% confidence intervals to account for small sample sizes and low detection frequencies. Logistic regression was used to assess the effects of harvest condition (pre-harvest vs post-harvest) and harvester section on the probability of prevalence. For processed lettuce harvester, pre-harvest samples and the conveyor belt section were selected as reference categories, representing baseline post-cleaning conditions and the primary product-contact surface, respectively. Model fitting included checks for sparse data and quasi-separation, and model outputs were interpreted cautiously when coefficient instability or wide confidence intervals were observed.

Microbial abundance was analyzed separately and restricted to DQ samples only, using log_10_- transformed surface-normalized concentrations (copies/cm^2^). In cases where DQ observations were too sparse or where quasi-separation across harvester sections limited reliable estimation, formal abundance models were not fit. Instead, abundance distributions were summarized descriptively. For reference, abundance of DQ+DNQ samples were visualized using boxplots with overlaid individual observations to illustrate variability (Fig S3 and S4). This strategy ensured that quantitative inference was restricted to reliably measurable concentrations while preserving sensitivity to low-level contamination events through prevalence-based analyses.

### 2.6. Portable On-farm LAMP to test harvester hygiene

We developed µPADs embedded with dried LAMP reagents (Wang et al., 2021). Using these µPADs, we performed LAMP assays on 96 harvester swab samples, directly using each swab resuspension to rehydrate the µPADs. Our device integrates a portable heater-imager system designed to facilitate LAMP testing. We refer to this complete unit as ‘FARM-LAMP’. To test the harvester hygiene on farms, LAMP reaction mixtures and 100 µPADs strips were fabricated in the laboratory (Wang et al., 2024a). Each testing strip had two µPAD: one with no primers and one with primers (Fig. S5). 27 mL of LAMP reaction mixtures were dried on each µPADs. The fabricated devices were packed in an insulated box with ice packs and transported to the testing site. All assays were conducted on-site using the FARM-LAMP system, which was powered by a Jackery Portable Power Station (500 W, 110 V) (Jackery, Explorer 500). For each sample, 27 μL of the swab resuspension was used to rehydrate µPADs using a micropipette (Eppendorf, USA). Both the µPADs were rehydrated with the samples. The rehydrated µPADs were then enclosed in reclosable polypropylene bags (Uline, S-17954) and incubated at 65°C for 60 minutes within the FARM-LAMP device. The imaging system captured time-lapse photos of the µPADs at one-minute intervals throughout the incubation period. Details regarding the heater fabrication and imaging system are provided in our previous manuscript (Wang et al., 2024a). The remaining swab resuspension samples were stored in 1.5 mL vials, kept on ice, and transported back to Purdue University, West Lafayette in a cooler box with ice packs via FedEx Priority Overnight for further analysis.

### 2.7. Image analysis and quantification for on-farm LAMP

Time-lapse images acquired through the FARM-LAMP system were analyzed to extract colorimetric data, with image processing conducted using the OpenCV library (code for analysis at Wang et al., 2024b). Each pixel was classified into weighted color bins, enabling the temporal quantification of sample positivity percentages. To enhance this analysis, derivative-based approaches were applied to assess variations in positivity rates over time, facilitating the estimation of DNA concentrations within the samples (Fig. S6). The LoD was determined, and the time-to-peak of the second derivative was plotted against *Bacteroidales* DNA concentrations on a logarithmic scale. A regression model was subsequently developed to quantify reaction concentrations based on the time-to-peak metric. The resulting regression equation was then used to calculate the concentration of field-tested samples by plugging in their respective time-to-peak values from the second derivative (Wang et al., 2024a).

## 3. Results

### 3.1. Establishing baseline levels of *Bacteroidales* on harvester surfaces using qPCR

We collected 400 swab samples from the surfaces of lettuce harvesters actively used during the season. We sampled two types of harvesters: (1) Processed lettuce harvesters, where the lettuce is collected and processed at a separate facility, and (2) Fresh-pack harvesters, where the lettuce is packed immediately on the farm post-harvest. The harvester equipment varied in architecture, and we swabbed the surfaces that directly contacted the lettuce during harvesting (i.e., zone 1 surfaces). After shipping, we resuspended the swabs in 500 µL of water and performed qPCR to measure the levels of the fecal contamination biomarker *Bacteroidales*. We derived the Cq values from the qPCR results and calculated copies/cm^2^ using a log-linear fit (Wang et al., 2024c).

qPCR data from harvester surface swabs contained a high proportion of ND results and few DQ observations. Accordingly, contamination was interpreted using a two-part framework that considered prevalence and abundance separately. Because DQ observations were sparse, inference was driven primarily by prevalence, whereas abundance among detected samples was summarized descriptively. Sample classification into ND, DNQ, and DQ followed the LoD (1 copy/reaction;5 copies/cm^2^;0.699 log_10_[copies/cm^2^];Cq ≤ 35.53) and LoQ (5 copies/reaction;25 copies/cm^2^;1.4 log_10_[copies/cm^2^];Cq ≤ 32.20) thresholds described in methods section 2.4 and 2.5.

For processed lettuce harvesters, we swabbed the conveyor belt, conveyor belt wall, curtain, tunnel, elevator, and funnel surfaces (an approximate area of 10 × 10 cm^2^) (Fig. S1). The sampling sites were selected based on how lettuce meets the harvester surfaces: from the ground to the collection box. We collected 204 samples for this type of harvester.

We observed a decreasing trend in the prevalence of *Bacteroidales* contamination across harvester sections, from the conveyor belt toward the funnel, corresponding to the increasing distance of lettuce from the ground and progression through the harvester (Fig. 1, Table S2). Overall prevalence was low and it declined from approximately 12.5% at the conveyor belt to 0% at the funnel (Table S2, average of pre-harvest and post-harvest conditions). Except for elevator (13.3% pre-harvest and 0% in post-harvest), prevalence was higher post-harvest for each section, particularly on upstream contact surfaces such as the conveyor belt (4.2% to 20.8%), conveyor belt wall (8.3% to 13.8%) and curtain (8.3% to 16.7%) (Fig. 1). Logistic regression indicated approximately two-fold higher odds of prevalence post-harvest; however, this effect did not reach statistical significance (p = 0.19, Table S3), likely due to sparse positive observations and quasi-separation across harvester sections.

**Fig. 1.**
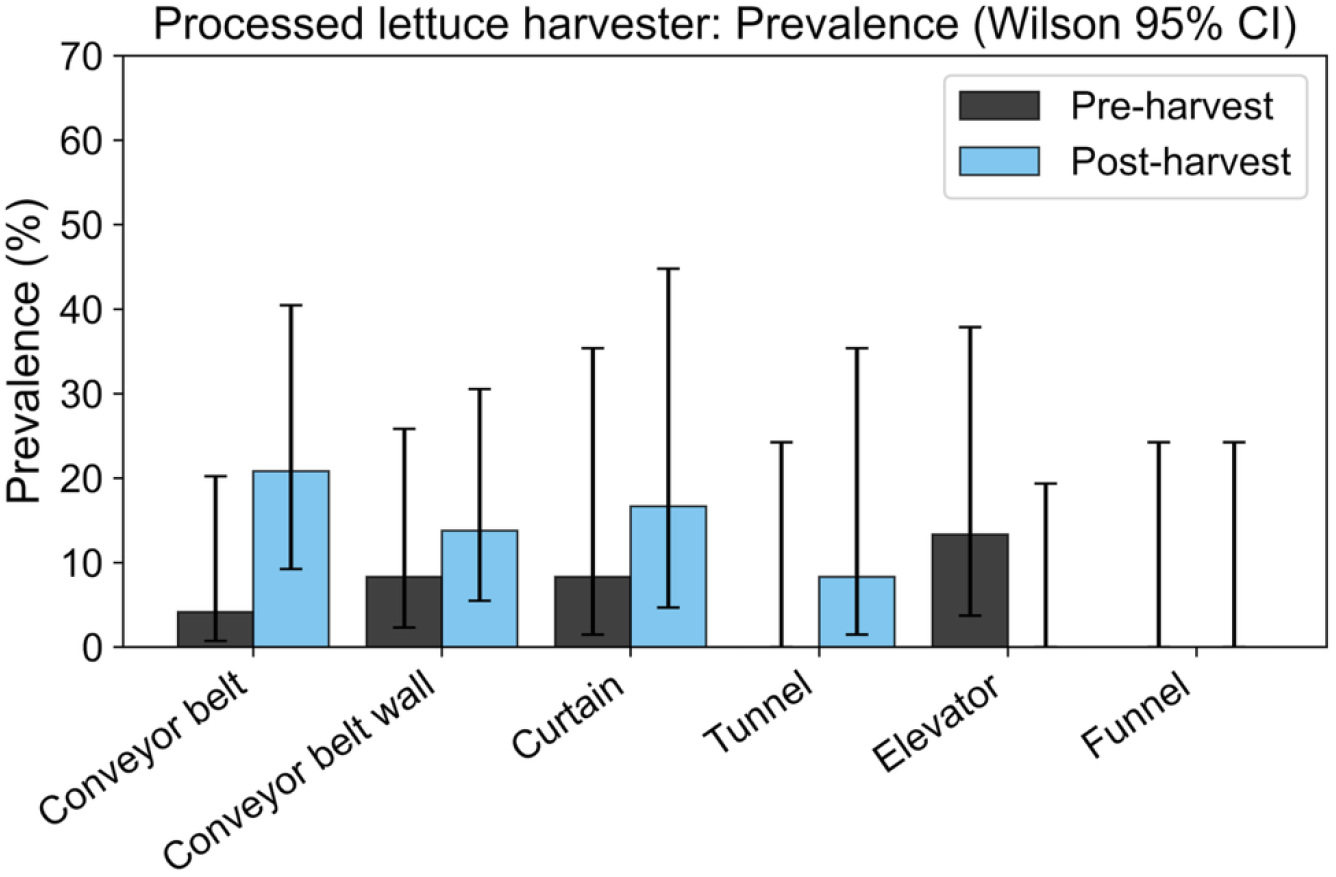
Prevalence (%) of *Bacteroidales* DNA on processed lettuce harvester surfaces by section and harvest stage based on qPCR data. Prevalence represents the proportion of samples classified as positive (DQ + DNQ) divided by all tested samples for each harvester section before (pre-harvest) and after (post- harvest) harvesting. Error bars indicate Wilson 95% confidence intervals.

DQ observations were rare, limiting the feasibility of statistical modeling of microbial abundance. As a result, contamination dynamics were interpreted primarily in terms of prevalence rather than abundance. Abundance analysis including all detected samples (excluding ND) is presented in the Supplementary Information (Fig. S3). Overall, *Bacteroidales* levels on processed lettuce harvesters were low (< 25 copies/cm^2^), with only a single sample reaching the DQ range (93 copies/cm^2^) (Fig. S3), indicating that harvest and processing operations primarily influence the probability of low-level contamination rather than increasing microbial loads once contamination is present. Elevated Cq values among detected samples likely reflect low target concentrations near the assay LoD and/or reduced efficiency in amplification in environmental matrices. Notably, the calibration curve generated from purified DNA yielded an amplification efficiency of 84.4%, and matrix-associated inhibition could further depress the efficiency and increase Cq in field samples. However, without confirmation using independent methods (e.g., culture-based or alternative molecular assays), it was not possible to distinguish true low-level contamination from potential false-positive detections near the LoD (Bustin et al., 2025).

For the fresh-pack lettuce harvester, a 10 × 10 cm^2^ area of the packing table was swabbed before and after harvesting (Fig. S2). The packing tables were covered with plastic linings, and all samples were collected directly from the plastic surface. In total, 196 samples were collected from this harvester type. Prevalence of the *Bacteroidales* marker was rare overall (Fig. 2, Table S4). No pre-harvest samples were positive, while a small proportion of post-harvest samples showed detectable signal (∼9%); however, wide Wilson 95% confidence intervals indicate uncertainty due to the limited number of positive observations. Logistic regression analysis suggested higher odds of detection post-harvest compared to pre-harvest; however, because no detections were observed in pre-harvest samples, odds ratios could not be estimated, and coefficient estimates were unstable due to quasi-separation (Table S5). Accordingly, results are interpreted based on observed prevalence rather than model-derived odds. Like the processed lettuce harvester, only a single DQ observation was recorded for the fresh-pack harvester, occurring in a post-harvest sample with an estimated concentration of 40 copies/cm^2^ (Fig. S4). This limited number of DQ observations precluded abundance modeling. Collectively, these results indicate that while detectable contamination was occasionally observed after harvest, quantifiable contamination on fresh-pack harvester surfaces was rare and highly sporadic. Overall, *Bacteroidales* levels on fresh-pack harvesters were low (<25 copies/cm^2^).

**Fig. 2.**
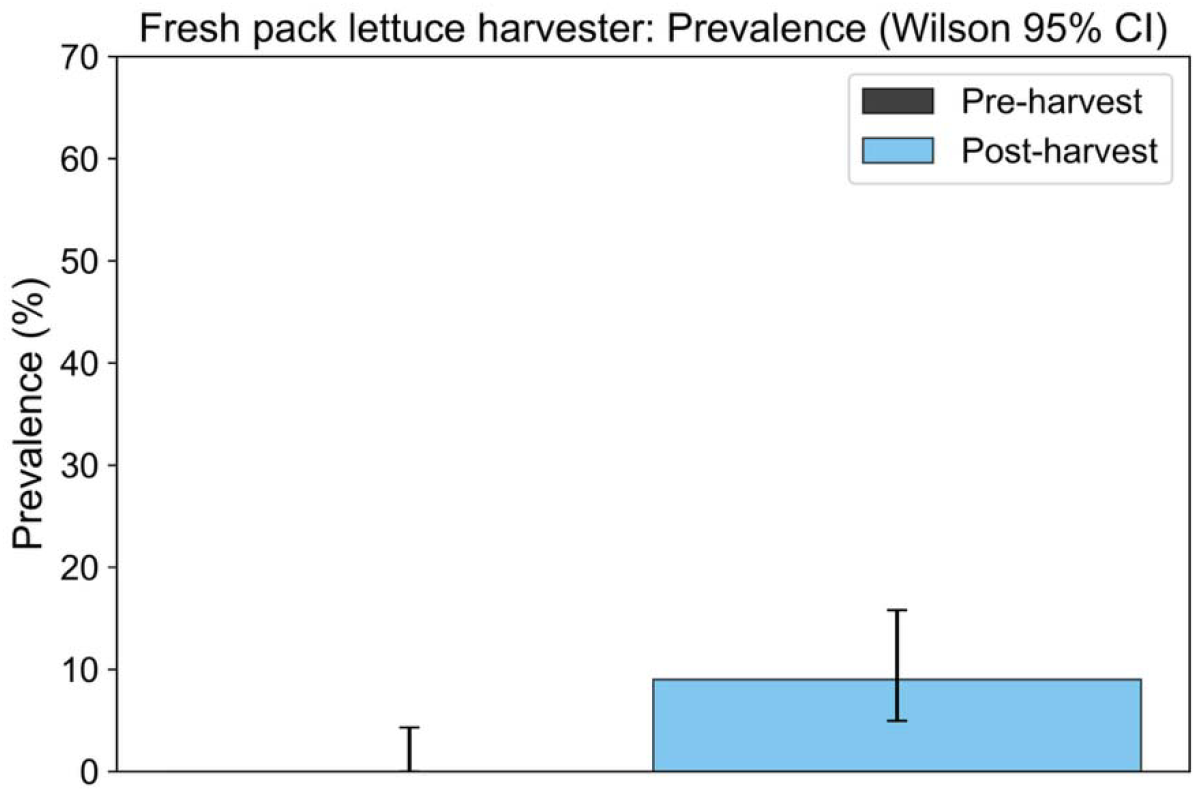
Prevalence (%) of the *Bacteroidales* DNA on fresh pack harvester surfaces by harvest stage based on qPCR data. Prevalence represents the proportion of samples classified as positive (DQ + DNQ) divided by all tested samples for before (pre-harvest) and after (post-harvest) harvesting. Error bars indicate Wilson 95% confidence intervals.

### 3.2. FARM-LAMP deployed on fresh-produce farms to test *Bacteroidales* on harvester surfaces

We deployed FARM-LAMP, a portable colorimetric LAMP testing platform integrating heating, imaging, and µPADs, for evaluating harvester hygiene. We validated the FARM-LAMP platform under real-world field conditions by performing assays directly in a vehicle-based mobile laboratory on farms. The platform’s effectiveness in detecting *Bacteroidales* on environmental samples, collection flags left on a farm for seven days was demonstrated in an earlier study by our group (Wang et al., 2024a).

We fabricated µPADs strips containing dried LAMP reagents (Wang et al., 2021). Each strip consisted of two separate µPADs: one included both LAMP reagents and primers, while the other contained LAMP reagents without primers (No Primer Control, NPC). We tested 96 harvester swab samples using FARM-LAMP, directly using each swab resuspension to rehydrate the µPADs. A volume of 27 µL of the resuspension was applied to each µPAD using a commercial micropipette (Eppendorf, USA). The rehydrated µPADs were sealed in resealable polypropylene bags (Uline) and heated at 65°C for 60 minutes in a portable heater. We included two negative controls (27 µL of nuclease-free water, No Template Control, NTC) and two positive controls (27 µL of *Bacteroides fragilis* DNA, 1 ng/reaction) with the samples. The µPADs are red due to the presence of the pH indicator phenol red in the reaction (Fig. 3A, Fig. S5). Upon amplification, the pH decreases, causing the color to change to yellow.

**Fig. 3.**
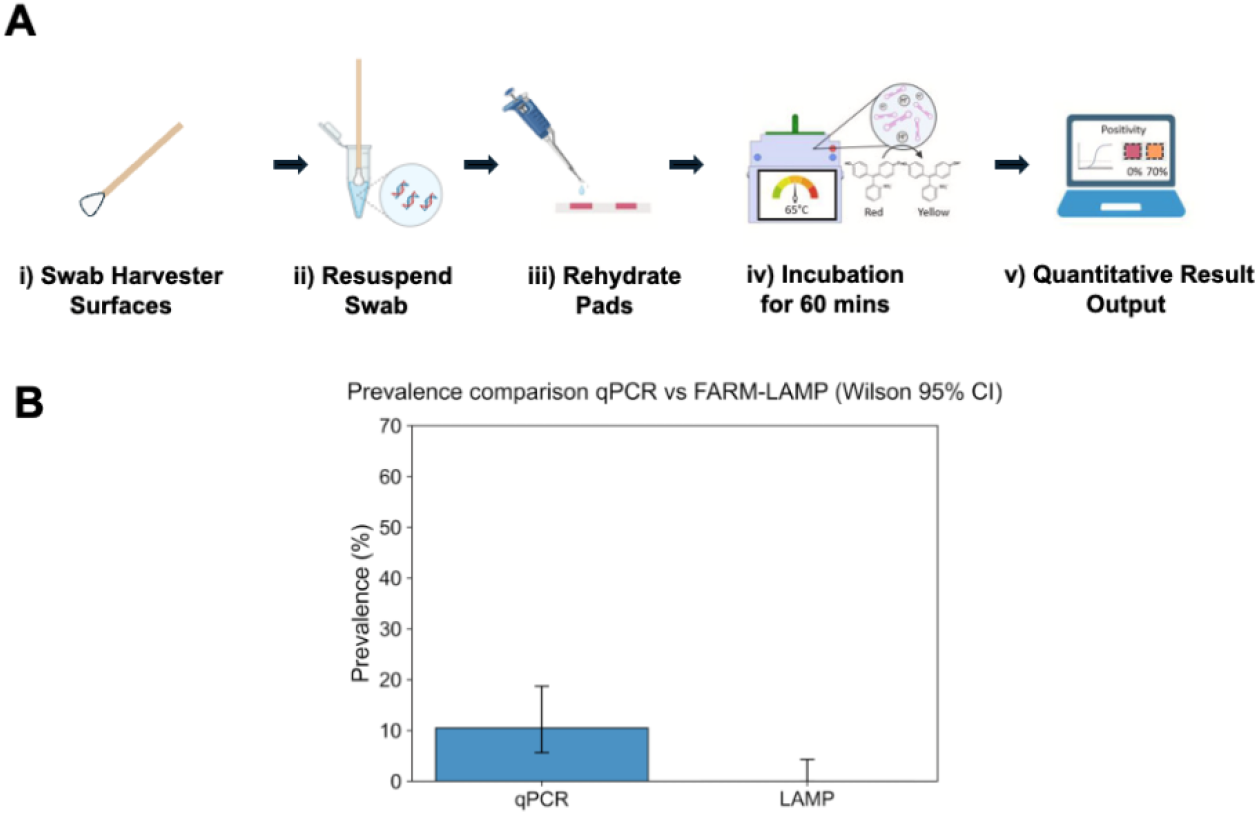
A) <u>Steps to test the hygiene of the harvester surface on farm:</u> i) Swab the surface of the harvester with a sterile swab. ii) Resuspend the swab in 500 μL of water. iii) Dispense the resuspension onto μPADs using a pipette. iv) Incubate the μPADs at 65°C for an hour, allowing LAMP amplification and a pH-induced color change from red to yellow in the presence of target DNA. v) An algorithm analyzes the color change of each μPAD to provide a quantitative readout (e.g., sample concentration in copies/μL). Figure adapted from Wang *et al* (Wang et al., 2024a). B) <u>Prevalence comparison between qPCR and FARM-LAMP</u>: Bars show the percentage of samples detected as positive by each molecular assay, with error bars representing Wilson 95% confidence intervals.

During the heating process, the imaging equipment captured time-lapse images of the µPADs every minute (Fig. S5). The imaging data were analyzed using a method previously detailed by Wang *et al* (Wang et al., 2024a). Each pixel in the image was classified into weighted bins to calculate sample positivity over time. We plotted the positivity percentages (Fig. S6 A) to visualize the quantitative color change and normalized the data by calculating the first and second derivatives to determine the onset of reaction pad amplification (Fig. S6 B). The limit of detection of FARM-LAMP assay is 100 copies/reaction (or 18.5 copies/cm^2^) (Wang et al., 2024a). Comparing the mean y-values of the first derivative to the lowest analyte concentration signal (0.75% positivity/min from 100 copies/reaction) reveals that all field samples and negative controls fell within this signal level, indicating a flat amplification curve (Fig. S6, A and B) (Wang et al., 2024a). Positive controls show a mean y-value of 1.34 % positivity/min, exceeding the LoD and confirming their positivity (mean positivity percentage 90.26%) (Fig. S6, C and D). The swab samples display a red color (negative) at the end of 60 minutes of heating (Fig. S5).

To evaluate the performance of FARM-LAMP relative to a laboratory-based molecular method, the same set of field samples was analyzed in parallel using qPCR (Fig. 3B). Detection outcomes were compared at the assay level using assay-specific LoD, and prevalence analysis. Overall, qPCR detected *Bacteroidales* in a small subset of samples (∼10% prevalence), whereas FARM-LAMP yielded no detections for this sample set (Fig. 3B). Paired detection outcomes between qPCR and FARM-LAMP were evaluated for samples tested using both assays (Table S6). Based on established LoD difference between the methods (qPCR: 1 copy/reaction, 5 copies/cm^2^ or Cq ≤ 35.53; FARM-LAMP: 100 copies/reaction, 18.5 copies/cm^2^ or time-to-peak second derivative ≤ 50.79 min) (Wang et al., 2024a), qPCR detected the target in a greater proportion of samples than FARM-LAMP. Specifically, among 96 paired samples, 10 were detected by qPCR but not by FARM-LAMP, no samples were detected by FARM-LAMP in the absence of qPCR detection, and the remaining 86 samples were negative by both assays (Table S6). This asymmetric detection pattern reflects the better analytical sensitivity of qPCR. A paired McNemar exact test confirmed a statistically significant difference in detection outcomes between the two methods (p = 0.002), driven entirely by discordant qPCR-positive/FARM-LAMP-negative results. Importantly, these discordant samples corresponded to low-level detections below the FARM-LAMP LoD, indicating that FARM-LAMP detection outcomes were consistent with its LoD, while avoiding false-positive detections. Consistent with this interpretation, our previous work demonstrated the utility of FARM-LAMP for detecting higher-burden positive field samples, supporting its application as a rapid, field-deployable screening tool when contamination levels exceed its LoD (Wang et al., 2024a).

## 4. Discussion

### 4.1. Baseline levels of *Bacteroidales* on lettuce harvester surfaces

We swabbed zone 1 harvester surfaces before harvesting (pre-harvest, following cleaning and sanitation [C/S]) and after harvesting (post-harvest, prior to C/S). Across both processed and fresh-pack harvester types, *Bacteroidales* concentrations were dominated by NDs and low-level detections near the assay LoD, with values less than 25 copies/cm^2^. Only two samples (out of 400 total) exceeded LoQ, with a maximum observed concentration of 93 copies/cm^2^. These findings establish baseline *Bacteroidales* levels on lettuce harvester zone 1 surfaces under commercial production conditions where farms adhered to standard produce safety and sanitation practices.

Previously, we reported baseline *Bacteroidales* levels of 0 or <LoD to 2 copies/cm^2^ on fresh produce farms using collection flags (Wang et al., 2024c). In contrast, we also showed that *Bacteroidales* concentrations are 10^3^-10^4^ copies/cm^2^ in fields adjacent to animal feeding operations (Wang et al., 2023) (Table 1). These studies indicate that while *Bacteroidales* levels on harvester surfaces are modestly elevated relative to general farm environments, they remain 500 to 5,000-fold lower than levels observed in high-risk regions influenced by nearby animal operations. This gradient supports the interpretation that harvester-associated contamination reflects low-level environmental transfer rather than direct fecal loading.

**Table 1:** Baseline levels of *Bacteroidales*.

| Sample type | *Bacteroidales* levels (copies/cm<sup>2</sup>) | Study |
| --- | --- | --- |
| Collection flag (on fresh produce farms) | 0 or <LoD -2 | (Wang et al., 2024c) |
| Collection flag (fields around animal unit) | 10<sup>3</sup>-10<sup>4</sup> | (Wang et al., 2023) |
| Swabs of harvester surfaces | <25 | This study |

*Bacteriodales* abundance and prevalence may vary when the harvester design is different, and zone 1 surfaces present cleaning challenges. Surface material and condition also influence deviations from baseline measurements. With prolonged use, wear and tear gradually change the surface properties of harvesting equipment. The duration of use that causes these changes varies by equipment part, for example, belts and chains typically show wear after 50-100 hours, while bearings and bushings require fresh grease every 10-20 hours (Inox Lubricants, 2024). This evolution can impact bacterial adherence, accumulation, and the formation of biofilms on the harvesting equipment (Kleine et al., 2019). Studies document cases where cleaning practices require adjustment due to increasing difficulty in removing bacteria from worn surfaces (Verran et al., 2001).

For processed lettuce harvesters, total prevalence exhibited a downstream decrease from the conveyor belt toward the funnel (Table S2), consistent with increasing distance from the soil and reduced direct contact with field material. Detection events were more frequently observed post- harvest on upstream surfaces, particularly the conveyor belt (pre-harvest ∼4% and post-harvest 20%), suggesting that these components may represent critical control points for managing low-level contamination probability. Similarly, for fresh pack harvester, prevalence was higher post-harvest (0 to 9%) (Table S4). However, due to sparse detectable observations and quasi-separation across sections, these patterns should be interpreted as trends rather than statistically significant differences.

Several other indicators are used to assess the cleanliness of food contact surfaces, including ATP (Leaman et al., 2023), and proteins (Schmitt and Moerman, 2016). Detecting ATP provides a broad view of cleanliness and detecting both microbial and non-microbial contaminants. Protein testing specifically detects residual proteins left on surfaces after cleaning. It is particularly useful for identifying food residues, including allergens, and it does not directly indicate microbial contamination. In contrast, fecal contamination indicators provide more targeted information about the risk of enteric pathogens, which are responsible for foodborne illnesses.

### 4.2. FARM-LAMP deployed on fresh produce farms to test harvester hygiene

µPADs are inexpensive to fabricate and use, with materials such as cellulose paper (Dong et al., 2023). We deployed a fully integrated, portable FARM-LAMP testing platform (comprising heating, imaging, and a µPADs-based LAMP assay) on a commercial lettuce farm to enable on-site detection of *Bacteroidales* within 60 minutes. Powered by a portable power station and operating from the back of a vehicle, this platform represents the first application of a µPADs based LAMP system for harvester hygiene assessment in fresh produce farms. Our approach sets the stage for future NAATs to be integrated into routine harvesting practices.

Although no contamination was detected in the harvester swabs, all positive controls exhibited a color change within 60 minutes, while negative controls showed no reaction, validating the performance of the FARM-LAMP. To further demonstrate its field applicability, we have previously tested samples near an animal unit, where the platform successfully detected *Bacteroidales*, underscoring its potential for *in-situ* testing (Wang et al., 2024a).

LAMP offers a solution to qPCR-related challenges, such as the need for expensive, non-portable lab equipment that requires sending samples to a laboratory (Ahmed et al., 2025; Kamel et al., 2025; Rafiq and Verma, 2025; Ranjbaran et al., 2024; Raut et al., 2026, 2026; Wang et al., 2022). However, both qPCR and LAMP share a limitation that they detect DNA from dead bacteria (Kaur et al., 2025). Non-viability does not pose a challenge in this application because we determine the quantities of indicator organisms’ DNA, not the pathogen itself. If we detect *Bacteroidales* DNA (whether from dead or live bacteria) at a concerning level, we investigate further for potential pathogen risk.

## 5. Conclusions

This study achieves the following three advances: 1) It establishes baseline *Bacteroidales* levels on lettuce harvester zone 1 surfaces, which were dominated by NDs and low-level detections, with concentrations ranging <25 copies/cm^2^. 2) This work provides insights into assessing fecal contamination risk from *Bacteroidales* levels by comparing the levels on harvester surfaces (<25 copies/cm^2^), fresh produce farms (0 or <LoD to 2 copies/cm^2^), and near animal feeding operations (10^3^-10^4^ copies/cm^2^). This demonstrates its potential as an indicator for fecal contamination risks in the fresh produce industry. 3) We deployed a fully integrated FARM-LAMP testing platform that enables on-site detection of contamination within one hour (reaction run time), combining the specificity and sensitivity of microbial techniques with the speed of chemical-based assays. This demonstration highlights the advantages of µPADs-based LAMP approaches for field applications to enable scalable routine testing, offering an effective solution for rapid hygiene monitoring. This technology offers a promising avenue to ensure food safety, meet stringent regulatory requirements, and protect public health (Dai et al., 2023).

However, this work has two limitations. 1) The presence of enteric bacteria does not always correlate with foodborne pathogen contamination, limiting the use of *Bacteroidales* as a risk indicator (Wang et al., 2024c). 2) This study does not account for the condition of harvesting machines (e.g., age, maintenance) during data collection.

Future efforts should focus on correlating *Bacteroidales* levels with pathogens, integrating real-time data analysis in FARM-LAMP device and automated sample processing. This could further enhance contamination risk assessments and improve response strategies. Additionally, developing testing protocols that account for harvester machine condition and environmental variability will strengthen hygiene monitoring practices. Future investigations could also include integrating soil, water, and product sampling to provide a more comprehensive understanding of contamination dynamics across the production environment.

## 6. Supporting Information

Additional experimental details, including materials and methods; supporting figures and tables; and references for supporting data.

## 7. Data Availability Statement

The data sets generated and/or analyzed during the current study are available in the Mendeley data repository at DOI: 10.17632/x24858cjvp.1

## 8. Declaration of AI and AI-assisted Technologies in the Writing Process

During the preparation of this work the authors used ChatGPT (https://chat.openai.com/) to check for grammar errors and improve the academic writing language. After using this tool/service, the authors reviewed and edited the content as needed. The authors take full responsibility for the content of the publication.

## 9. Credit Author Statement

**Simerdeep Kaur:** Conceptualization, Data curation, Formal analysis, Investigation, Methodology, Software, Validation, Visualization, Writing – original draft, Writing – review & editing. **Jiangshan Wang:** Conceptualization, Data curation, Investigation, Methodology, Software, Writing – review & editing. **Ashley Kayabasi:** Methodology, Software, Writing – review & editing. **Ishaan Rath:** Methodology, Writing – review & editing. **Ilan Benschikovski:** Methodology, Writing – review & editing. **Bibek Raut:** Investigation, Validation, Writing – review & editing. **Kyungyeon Ra:** Investigation, Validation, Writing – review & editing. **Mohit S. Verma:** Conceptualization, Funding acquisition, Methodology, Project administration, Supervision, Writing – review & editing.

## 10. Funding Statement

This work was funded in part by the Center for Produce Safety (CPS Award Number: 2021CPS12), the California Department of Food and Agriculture (CDFA Agreement No. 20- 0001-054-SF), and the U.S. Department of Agriculture’s (USDA) Agricultural Marketing Service (USDA Cooperative Agreement No. USDA-AMS-TM-SCBGP-G-20-0003). The project entitled “Field evaluation of microfluidic paper-based analytical devices for microbial source tracking” was funded in whole or in part through a subrecipient grant awarded The Center for Produce Safety through the California Department of Food and Agriculture (2020) Specialty Crop Block Grant Program and the U.S. Department of Agriculture’s (USDA) Agricultural Marketing Service. We acknowledge support from The Center for Produce Safety (2023CPS11) and the U.S. Department of Agriculture’s (USDA) Agricultural Marketing Service through grant AM22SCBPCA1133 for “Testbeds for microbial source tracking using microfluidic paper-based analytical devices”. Any opinions, findings, conclusions, or recommendations expressed in this publication or audiovisual are those of the author(s) and do not necessarily reflect the views of The Center for Produce Safety, the California Department of Food and Agriculture, or the U.S. Department of Agriculture (USDA).

We are grateful for the collaboration with Patricia Contreras, Felice Arboisiere, and Natalie Dyenson from Dole Fresh Vegetables, Inc. and Dole Food Company, Inc. for their assistance in collecting samples in the field.

