## Supporting Information for "*Bacteroidales* on Harvesters: Baseline Prevalence and Abundance"

**For**

**SUPPORTING TABLES**

| **Primer** | **Sequence (5’ - 3’)** | **Reference** |
| --- | --- | --- |
| Universal.*Bacteroidales*.16S rRNA.1.F3 | TGCGGGTATCGAACAGGATT | (Wang et al., 2023) FARM-LAMP |
| Universal.*Bacteroidales*.16S rRNA.1.B3 | GGTAAGGTTCCTCGCGTATC |  |
| Universal.*Bacteroidales*.16S rRNA.1.FIP | TTAACGCTTTCGCTTGGCCACAGTAGTCCGCACGGTAAACG |  |
| Universal.*Bacteroidales*.16S rRNA.1.BIP | GTACGCCGGCAACGGTGAAAACATGTTCCTCCGCTTGTG |  |
| Universal.*Bacteroidales*.16S rRNA.1.LF | GGCCGAACAGCGAGCAT |  |
| Universal.*Bacteroidales*.16S rRNA.1.LB | CAAAGGAATTGACGGGGGC |  |
| GenBac3.FP | GGGGTTCTGAGAGGAAGGT | (Dick and Field, 2004; Siefring et al., 2008) qPCR |
| GenBac3.RP | CCGTCATCCTTCACGCTACT |  |
| GenBac3.Probe | /FAM/CAATATTCC/ZEN/TCACTGCTGCCTCCCGTA/IABkFQ/ |  |

Table S1: FARM-LAMP and qPCR primers

Table S2: Prevalence of the target on processed lettuce harvester surfaces by section and harvest condition. Prevalence was defined as (quantifiable detection [DQ + DNQ]) divided by the number of samples tested with Wilson 95% confidence intervals.

| Section | Condition | N tested | N detected | Prevalence (%) | 95% CI (lower limit) | 95% CI (upper limit) | Total prevalence (%) |
| --- | --- | --- | --- | --- | --- | --- | --- |
| Conveyer belt | Pre-harvest | 24 | 1 | 4.2 | 0.007 | 0.202 | 12.5 |
| Conveyer belt | Post-harvest | 24 | 5 | 20.8 | 0.092 | 0.405 |  |
| Conveyer belt wall | Pre-harvest | 24 | 2 | 8.3 | 0.023 | 0.258 | 11.3 |
| Conveyer belt wall | Post-harvest | 29 | 4 | 13.8 | 0.055 | 0.305 |  |
| Curtain | Pre-harvest | 12 | 2 | 8.3 | 0.015 | 0.354 | 12.5 |
| Curtain | Post-harvest | 12 | 1 | 16.7 | 0.047 | 0.448 |  |
| Tunnel | Pre-harvest | 12 | 0 | 0 | 0 | 0.242 | 4 |
| Tunnel | Post-harvest | 12 | 1 | 8.3 | 0.015 | 0.354 |  |
| Elevator | Pre-harvest | 15 | 2 | 13.3 | 0.379 | 0.379 | 6.5 |
| Elevator | Post-harvest | 16 | 0 | 0 | 0 | 0.242 |  |
| Funnel | Pre-harvest | 12 | 0 | 0 | 0 | 0.242 | 0 |
| Funnel | Post-harvest | 12 | 0 | 0 | 0 | 0.242 |  |

Table S3: Logistic regression results for contamination prevalence on the processed lettuce harvester. Pre-harvest samples and the conveyor belt section were selected as reference categories because they represent the baseline condition and primary product-contact surface, respectively.

| **Predictor** | **Coefficient (β)** | **Std. Error** | **z value** | **p value** | **95% CI (lower limit)** | **95% CI (upper limit)** |
| --- | --- | --- | --- | --- | --- | --- |
| Intercept | −2.3406 | 0.550 | −4.252 | <0.001 | −3.419 | −1.262 |
| Post-harvest | 0.6980 | 0.527 | 1.324 | 0.186 | −0.335 | 1.731 |
| Conveyor belt wall | −0.1457 | 0.619 | −0.235 | 0.814 | −1.360 | 1.068 |
| Curtain | 0.0005 | 0.761 | 0.001 | 1.000 | −1.491 | 1.491 |
| Tunnel | −1.1988 | 1.114 | 1.076 | 0.282 | −3.382 | 0.985 |
| Elevator | −0.7452 | 0.855 | −0.872 | 0.383 | −2.421 | 0.931 |
| Funnel | −11.2798 | 150.928 | −0.075 | 0.940 | −307.093 | 284.534 |
| Tunnel | −1.1988 | 1.114 | −1.076 | 0.282 | −3.382 | 0.985 |

Table S4: Prevalence of the target on fresh pack lettuce harvester surfaces pre- and post-harvest. Prevalence was defined as quantifiable detection (DQ + DNQ) divided by the number of samples tested, with Wilson 95% confidence intervals.

| **Condition** | **N tested** | **N detected** | **Prevalence (%)** | **95% CI (lower limit)** | **CI (higher limit)** |
| --- | --- | --- | --- | --- | --- |
| Pre-harvest | 85 | 0 | 0 | 0 | 0.043 |
| Post-harvest | 111 | 10 | 9 | 0.049 | 0.158 |

Table S5: Logistic regression results for contamination prevalence on fresh pack harvester. Pre-harvest samples were selected as reference categories because they represent the baseline condition, respectively.

| **Predictor** | **Coefficient (β)** | **Std. Error** | **z-value** | **p-value** | **95% CI (lower limit)** | **95% CI (upper limit)** |
| --- | --- | --- | --- | --- | --- | --- |
| Intercept | −14.12 | 126.13 | −0.11 | 0.911 | −261.32 | 233.09 |
| Condition (Post) | 11.80 | 126.13 | 0.09 | 0.925 | −235.40 | 259.01 |

Table S6: Truth table comparing detection outcomes between qPCR and FARM-LAMP for samples tested using both assays

| **Truth table** | | |
| --- | --- | --- |
|  | LAMP Detected (+) | LAMP Not Detected (–) |
| qPCR Detected (+) | 0 | 10 |
| qPCR Not Detected (–) | 0 | 86 |

**SUPPORTING FIGURES**

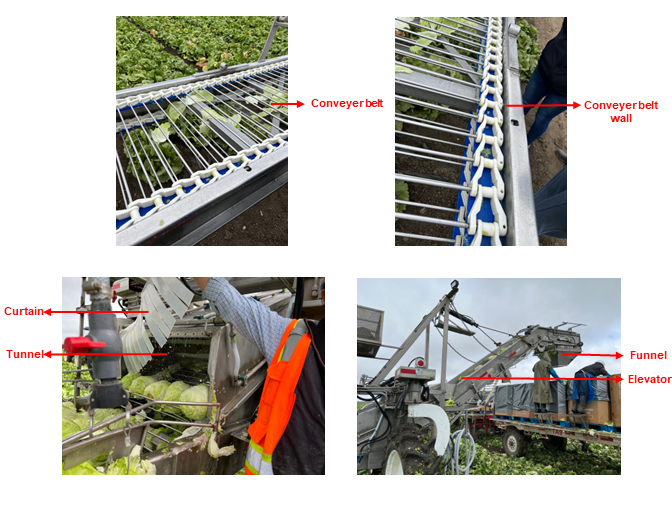

Figure S1: Swabbing sites for processed lettuce harvester

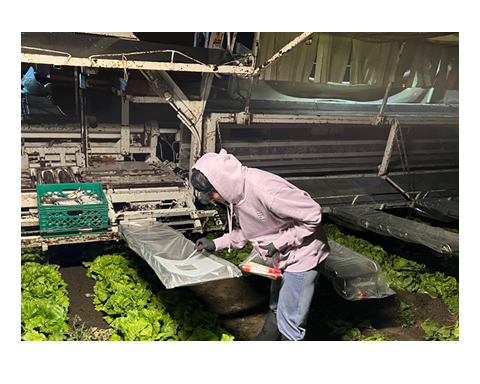

Figure S2: Swabbing site (packing table) for fresh pack lettuce harvester. Image is of the author (SK).

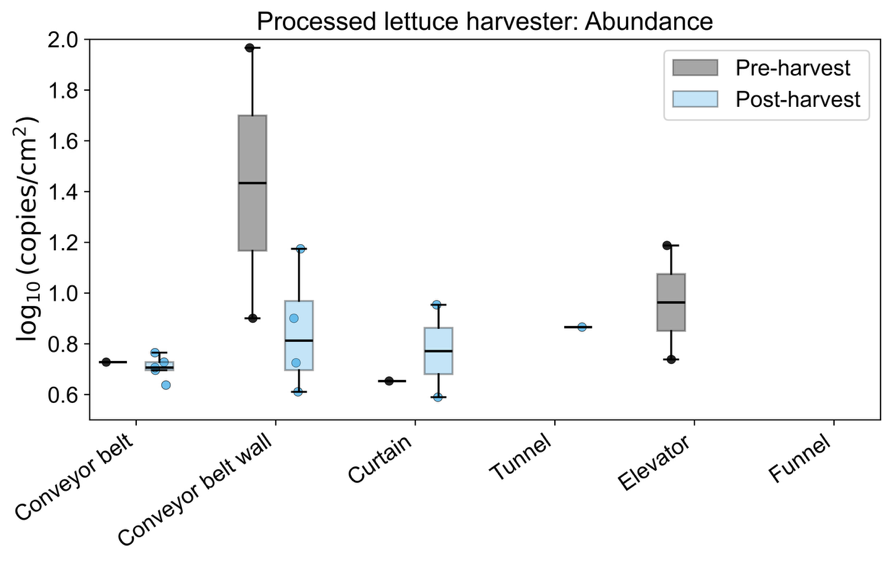

**Figure S3. Abundance of *Bacteroidales* on processed lettuce harvesters pre- and post-harvest operations for each section:** Boxplots show log_10_-transformed copies per cm^2^ (log_10_ copies/cm^2^). Individual points indicate measured samples; boxes denote the interquartile range (IQR), center lines indicate medians, and whiskers represent values within 1.5× IQR. All detected (DQ+DNQ) samples were included in abundance calculations, while non-detects were excluded. These plots illustrate differences in microbial load across equipment components and between operational stages.

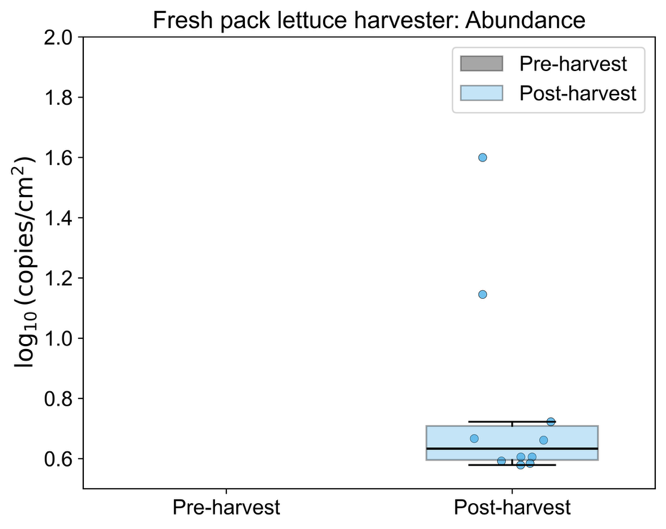

**Figure S4: Abundance of *Bacteroidales* on fresh pack lettuce harvesters pre- and post-harvest operations:**
Boxplots show log_10_-transformed copies per cm^2^ (log_10_ copies/cm^2^). Individual points indicate measured samples; boxes denote the interquartile range (IQR), center lines indicate medians, and whiskers represent values within 1.5× IQR. All detected (DQ+DNQ) samples were included in abundance calculations, while non-detects were excluded. These plots illustrate differences in microbial load across operational stages.

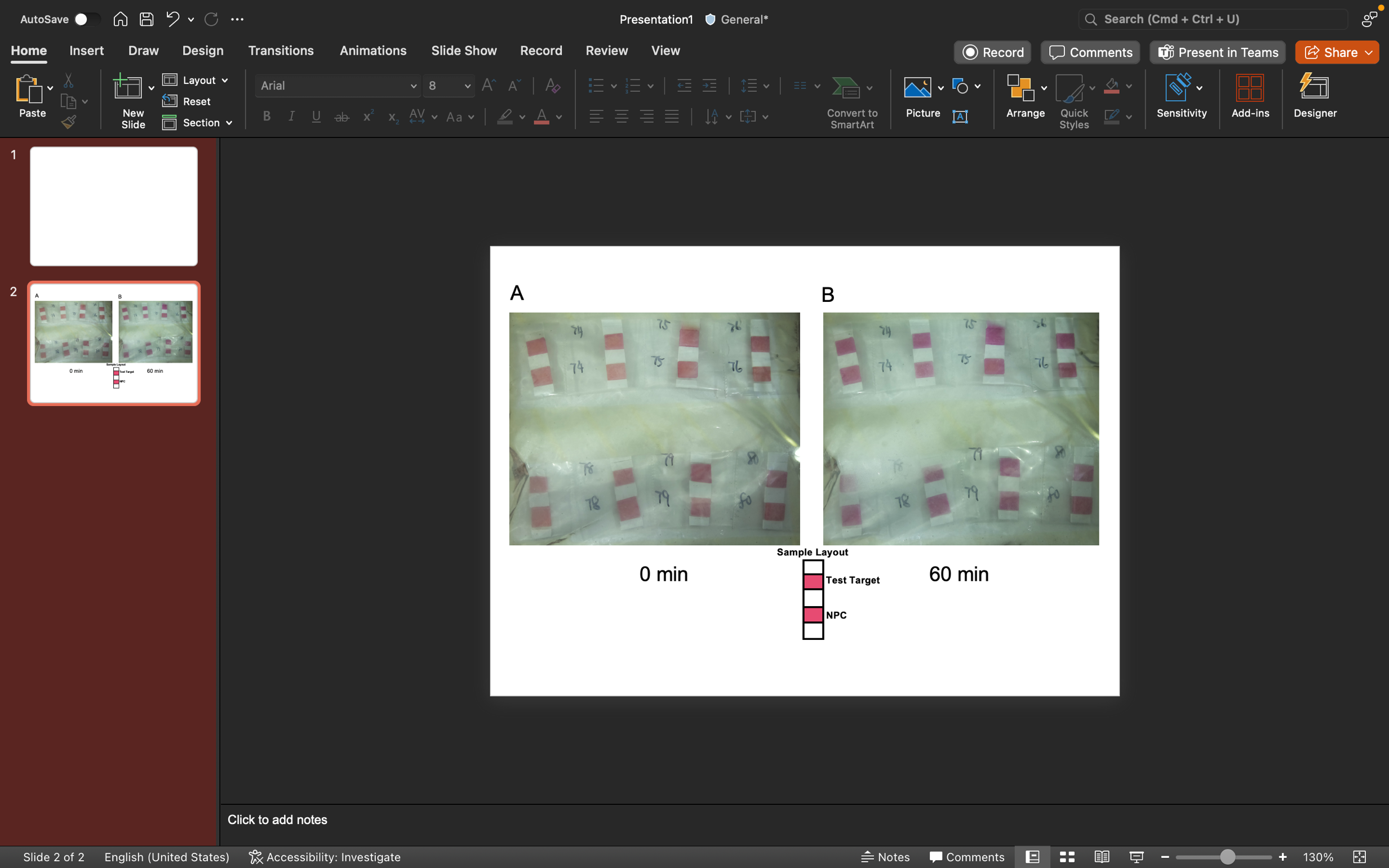

Figure S5: Images captured using the imaging system within the integrated heating and imaging setup. The assay was conducted for 60 minutes. (A) shows the image taken at 0 minutes, while (B) shows the image taken at 60 minutes. These images display μPADs labeled with their respective sample IDs, with their layout shown at the bottom. The upper μPAD represents the complete LAMP reaction, including primers, while the lower μPAD serves as the No Primer Control (NPC), which lacks primers and is used to validate each strip.

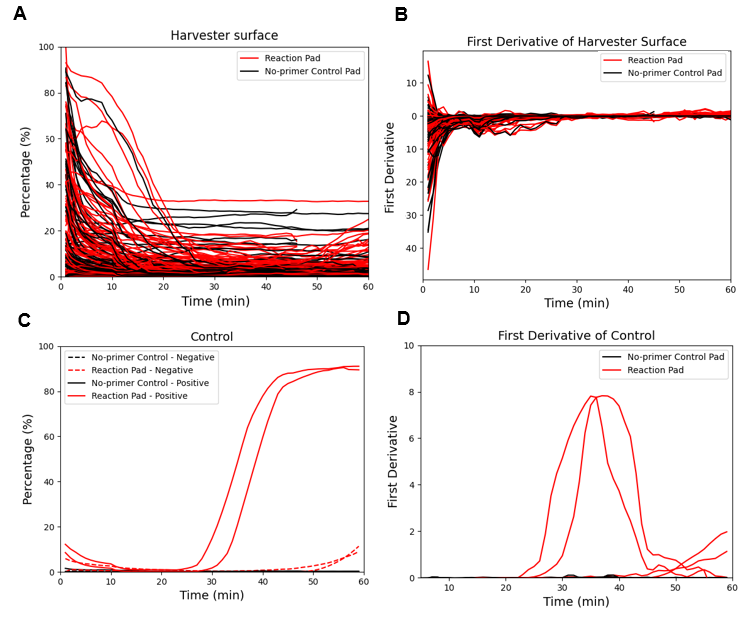

Figure S6: Image analysis FARM-LAMP: A) The percentage of positivity over time for each field sample. To visualize the color change qualitatively, the curves were smoothed using a moving average filter. B) First derivative of the positivity percentage for samples. C) The percentage of positivity over time for each control sample. Similar to field sample, a moving average filter was applied to smooth the curves and enable qualitative assessment of the color change. D) First derivative of the positivity percentage for controls.
